# Sumoylation regulates central spindle protein dynamics during chromosome segregation in oocytes

**DOI:** 10.1101/584763

**Authors:** Federico Pelisch, Laura Bel Borja, Ellis G. Jaffray, Ronald T. Hay

## Abstract

Meiotic spindles in most species lack centrosomes and the mechanisms that underlie faithful chromosome segregation in acentrosomal meiotic spindles are not well understood. In *C. elegans* oocytes, spindle microtubules exert a poleward force on chromosomes dependent on the microtubule-stabilising protein CLS-2^*CLASP*^. The kinase BUB-1^*Bub*1^ and CLS-2^*CLASP*^ localise in the central-spindle and display a dynamic localisation pattern throughout anaphase but the signals regulating their anaphase-specific localisation remains unknown. We have shown that SUMO regulates BUB-1 localisation during metaphase I. Here, we found that SUMO modification of BUB-1^*Bub*1^ is regulated by the SUMO E3 ligase GEI-17 and the SUMO protease ULP-1. SUMO is required for BUB-1 localisation in between segregating chromosomes during early anaphase I, and SUMO depletion partially phenocopies BUB-1^*Bub*1^ depletion. We also show that CLS-2^*CLASP*^ is subject to SUMO-mediated regulation. Overall, we provide evidence for a novel, SUMO-mediated control of protein dynamics during early anaphase I in oocytes.

## Introduction

Faithful chromosome partitioning is essential for accurate cell division and is achieved by physically separating chromatids or paired homologous chromosomes, in a process referred to as chromosome segregation. This is achieved by a complex and dynamic structure known as the spindle (Wittmann et al., 2001; Gadde and Heald, 2004; Dumont and Desai, 2012). Spindles consist of microtubules (MTs) and accessory proteins and spindle MTs are classified according to their location within the spindle and the structures they contact. Some MTs contact chromosomes through the kinetochore, a multi-protein complex that assembles on specific regions on chromosomes called centromeres (Tanaka and Desai, 2008; Verdaasdonk et al., 2011; Godek et al., 2015; Musacchio and Desai, 2017; Prosser and Pelletier, 2017). During anaphase, while chromosomes are segregating, MTs populate the interchromosomal region creating the central-spindle. While many studies of MT-dependent chromosome movement focused on pulling forces generated by kinetochore MTs (kMTs) making end-on contacts with chromosomes (Cheeseman, 2014), there is also evidence for pushing forces that are exerted on the segregating chromosomes (Khodjakov et al., 2004; Nahaboo et al., 2015; Laband et al., 2017; Vukušić et al., 2017; Yu et al., 2019).

The nematode *Caenorhabditis elegans* (*C. elegans*) contains holocentric chromosomes (Maddox et al., 2004) and has served as an extremely useful system to uncover mechanisms of meiosis and mitosis for almost twenty years (Oegema et al., 2001; Desai et al., 2003; Cheeseman et al., 2004; Cheeseman et al., 2005; Monen et al., 2005). Meiosis is a specialized cell division with two successive rounds of chromosome segregation that reduce the ploidy and generates these haploid gametes (Ohkura, 2015; Duro and Marston, 2015, Severson et al., 2016)‥ During meiosis I homologous chromosomes segregate while sister chromatid cohesion is maintained. During meiosis II, sister chromatid cohesion is lost, reminiscent of mitotic chromosome segregation (Dumont and Desai, 2012; Duro and Marston, 2015; Bennabi et al., 2016; Severson et al., 2016). During *C. elegans* female meiosis, kinetochores disassemble in early anaphase I and appear to be dispensable for chromosome segregation (Dumont et al., 2010; Hattersley et al., 2016; McNally et al., 2016). In addition, tomographic reconstruction in electron microscopy of the *C. elegans* female meiotic spindle revealed that during anaphase I, central-spindle MTs transition from a lateral to an end-on orientation (Laband et al., 2017; Redemann et al., 2018; Yu et al., 2019). Therefore, while the balance between central-spindle MT-and kMT-driven forces may vary in different spindles, the former seems most important during female meiosis in *C. elegans*. Many central-spindle proteins begin to concentrate between homologous chromosomes during prometaphase in a ring-shaped structure (hereafter ring domain), which marks the site of cohesion loss (Dumont et al., 2010; Muscat et al., 2015; Pelisch et al., 2017; Wignall and Villeneuve, 2009). The metaphase I ring domain consists of key cell division regulators including the chromosomal passenger complex (CPC) components AIR-2^*AuroraB*^, ICP-1^*INCENP*^, BIR-1^*Survivin*^ (Schumacher et al., 1998; Speliotes et al., 2000; Kaitna et al., 2002; Rogers et al., 2002; Romano et al., 2003), the checkpoint kinase BUB-1^*Bub*1^ (Monen et al., 2005; Dumont et al., 2010), Condensin I components (Collette et al., 2011), the kinesin KLP-7^*MCAK*^ (Connolly et al., 2015; Han et al., 2015.; Gigant et al., 2017), and the chromokinesin KLP-19^*Kif4A*^ (Wignall and Villeneuve, 2009). We recently showed that a number of these components are held together by a combination of covalent SUMO modification and non-covalent SUMO interactions (Pelisch et al., 2017). SUMO conjugation/localisation is highly dynamic during meiosis and the functional significance of this highly regulated SUMO modification in the ring domain composition once it is formed and throughout anaphase remains largely unexplored. Furthermore, the role of this ring domain itself during chromosome segregation has remained elusive. During anaphase the ring domain stretches and its composition changes rapidly, leading to the recruitment of SEP-1^*Separase*^ (Muscat et al., 2015), MDF-1^*Mad*1^ (Moyle et al., 2014), and CLS-2^*CLASP*^ (Dumont et al., 2010; Laband et al., 2017). CLS-2 exhibits a BUB-1-dependent kinetochore localisation until metaphase I and localises within the central spindle during anaphase. Additionally, BUB-1 and CPC components are also present within the central spindle. The limited evidence on the dynamics of these proteins during meiosis I suggests that they do not necessarily occupy the same domains throughout anaphase (Dumont et al., 2010; Davis-Roca et al., 2017; Mullen and Wignall, 2017; Davis-Roca et al., 2018). Considering that i) the CPC and CLS-2 are essential for chromosome segregation and ii) BUB-1 also plays a role during chromosome segregation, we sought to focus our attention on these proteins and characterise their dynamics during anaphase I. Given the relevance of the CPC, BUB-1, and CLS-2, and the established role for SUMO during metaphase, we sought to understand the mechanisms underlying these proteins’ localisations and interactions. We hypothesised that SUMO modification regulates these proteins’ dynamic localisations because i) ring domain proteins are SUMO substrates (Pelisch et al., 2014; Pelisch et al., 2017), ii) other ring components (i.e. GEI-17 and BUB-1) can interact non-covalently with SUMO (Pelisch et al., 2017), and iii) the reversible/dynamic nature of this post-translational modification (PTM) would allow for rapid changes in the protein interaction network within the meiotic spindle.

Here, we show that the key cell division regulators AIR-2, BUB-1, and CLS-2 exhibit highly dynamic localisation patterns during meiosis I. While AIR-2 and BUB-1 co-localise during early anaphase, these proteins subsequently occupy complementary domains as chromosomes segregate. Conversely, while reducing its co-localisation with BUB-1, AIR-2 co-localisation with CLS-2 increases as anaphase progresses. We found that the precise spatial and temporal localisation of these proteins is dependent on SUMO. We demonstrate that the SUMO modification status of BUB-1 is controlled by the SUMO E3 ligase GEI-17 and by the SUMO protease ULP-1. Overall, sumoylation is a key post-translational modification for the correct localisation of key proteins such as BUB-1 and CLS-2 during female meiosis.

## Results

### Dynamic localisation of SUMO and central spindle proteins during anaphase I

We previously showed that SUMO localises in the midbivalent ring domain and regulates KLP-19 and BUB-1 localisation during metaphase I (Pelisch et al., 2017). Based on these observations, we addressed the role of the SUMO conjugation pathway during meiotic chromosome segregation in *C. elegans* oocytes. During early anaphase, the midbivalent rings stretch into rod-like structures within the central spindle and SUMO remains strongly concentrated in these structures (Figure 1A). High resolution live imaging of dissected oocytes expressing GFP-tagged SUMO shows that the SUMO signal increases after anaphase onset, peaking during early anaphase (Figure 1A, B and Supplementary Movie 1). This is followed by a diffusion throughout the central spindle and a sharp decrease in intensity at 100 seconds after anaphase onset (Figure 1A, B). We have shown before that GEI-17, the sole *C. elegans* PIAS orthologue, is the key meiotic SUMO E3 ligase (Pelisch et al., 2017). The SUMO E3 ligase GEI-17 displays a localisation pattern similar to that of SUMO, Supporting the notion that SUMO conjugation is actively taking place during early anaphase (Figure 1C, D).

**Fig. 1.**
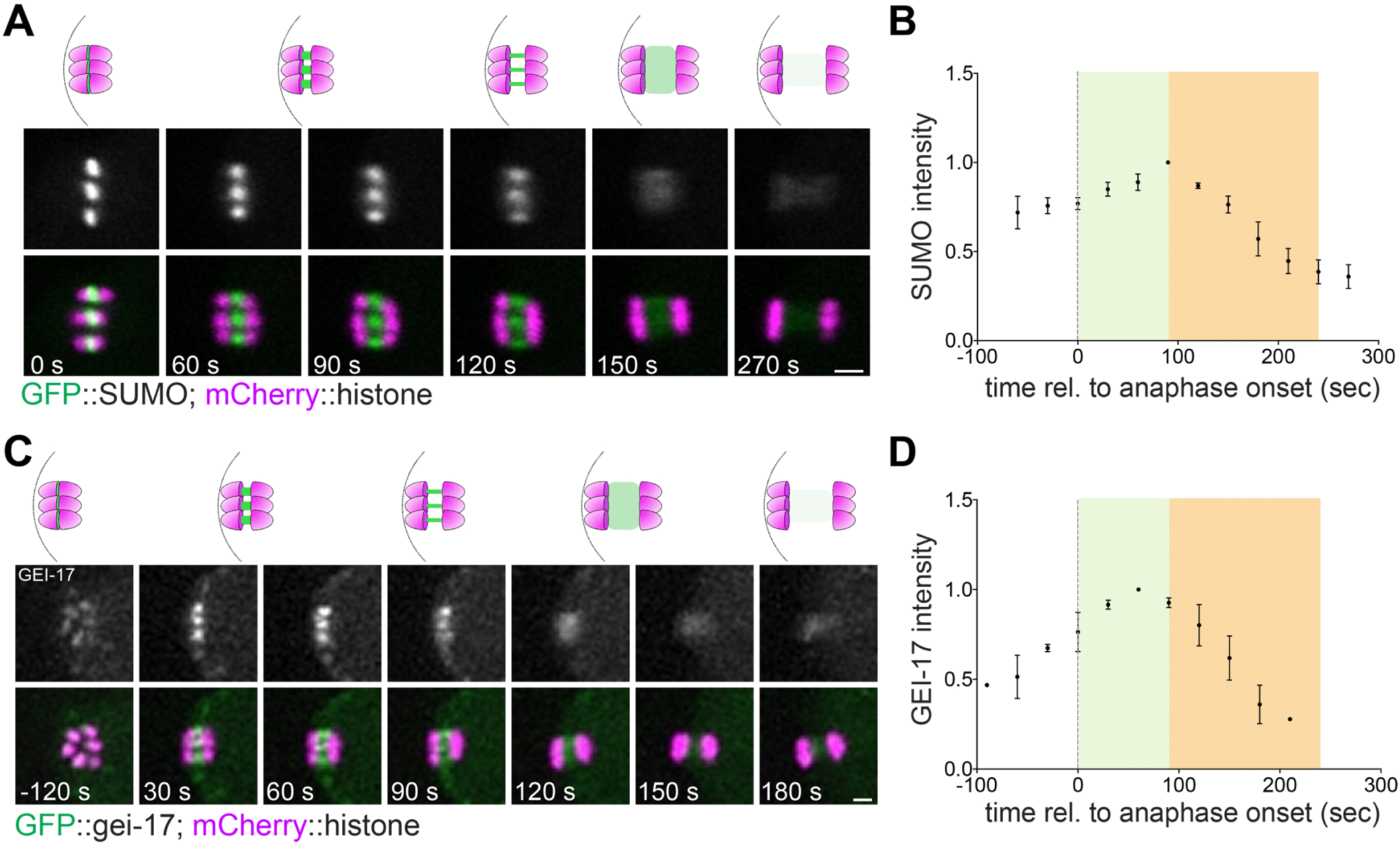
SUMO dynamics during anaphase I. **A.** SUMO localisation throughout meiosis I was followed in live oocytes from a strain expressing GFP::SUMO. A single z-slice is shown in the images. **B.** Quantitation of the SUMO signal from (A). **C.** The SUMO E3 ligase GEI-17 localisation was followed throughout meiosis I in oocytes expressing GFP::GEI-17. **D.** Quantitation of GFP::GEI-17 from (C). Scale bars, 2 µm.

To assess the role of SUMO during anaphase I progression, we investigated the localisation and role of two proteins shown to play key roles during meiotic chromosome segregation: BUB-1 and AIR-2 (Dumont et al., 2010; Kaitna et al., 2002; Rogers et al., 2002; Muscat et al., 2015; Laband et al., 2017). AIR-2 concentrates in the midbivalent ring domain (Kaitna et al., 2002; Rogers et al., 2002), while BUB-1 is present in the ring domain and also in kinetochores (Monen et al., 2005; Dumont et al., 2010; Laband et al., 2017). In agreement with this, we observed a strong AIR-2 and BUB-1 co-localisation in the midbivalent ring (Figure 2A, cyan arrows). During anaphase, BUB-1 remains at the core of the ring domains, while AIR-2 concentrates more on the edges of the rod-like structures, closer to chromosomes (Figure 2A, yellow arrows). Later in anaphase (judged by the chromosome separation), AIR-2 and BUB-1 occupy completely non-overlapping domains within segregating chromosomes (Figure 2A). During late anaphase, BUB-1 signal is lost while AIR-2 concentrates solely in the central spindle, where microtubules (not shown in the figure) have populated the entire area (Figure 2A). Such AIR-2 and BUB-1 dynamic localisations were confirmed by live imaging of dissected oocytes (Figure 2B). We noted that the strong BUB-1/AIR-2 co-localisation occurs during metaphase and early anaphase, coinciding with the peak in SUMO conjugation. We therefore compared SUMO localisation to that of BUB-1 and AIR-2 in live oocytes.

**Fig. 2.**
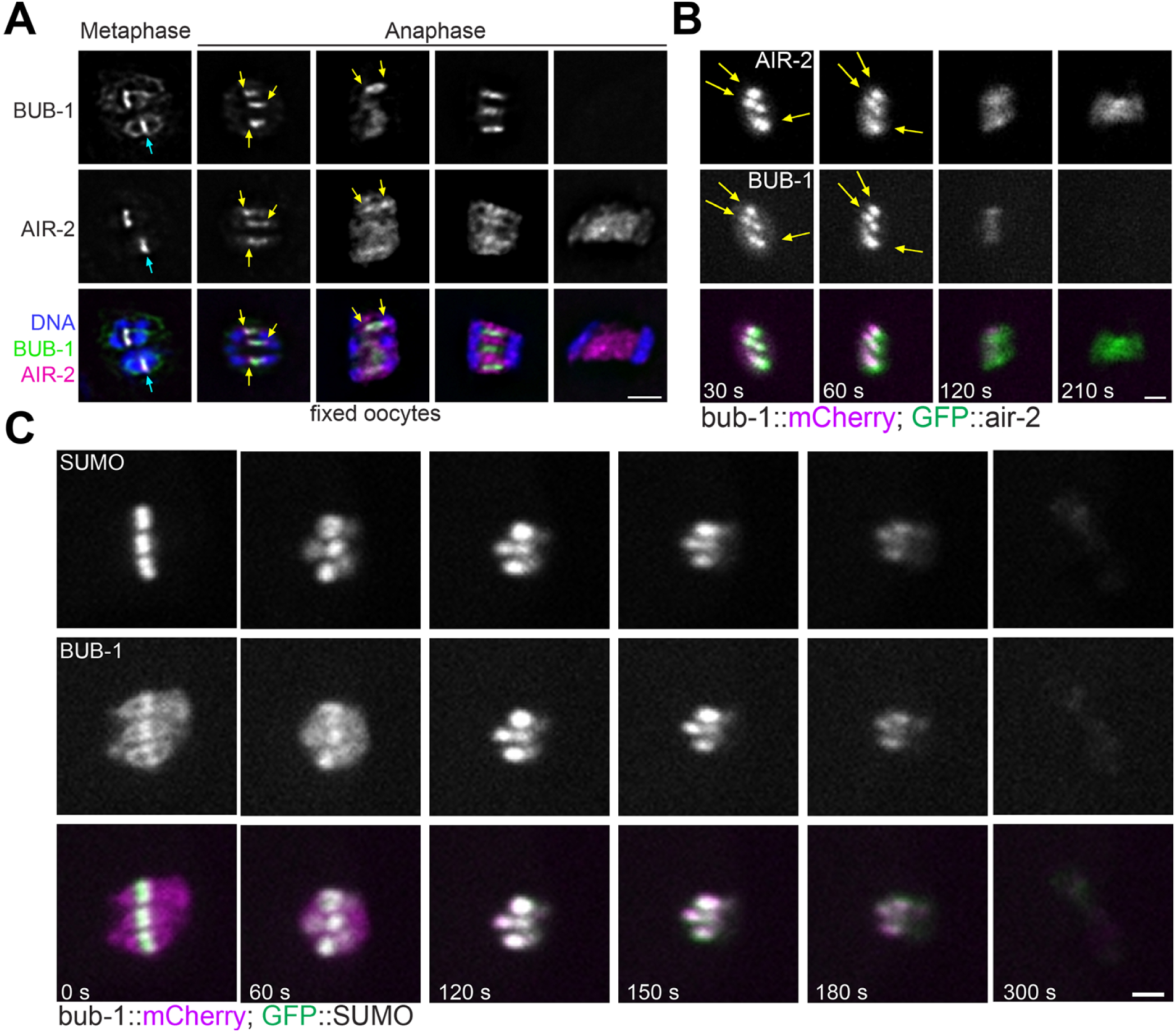
SUMO, BUB-1, and AIR-2 dynamics during anaphase I. **A.** BUB-1 and AIR-2 localisation at different meiosis I stages were analysed in fixed samples. Note that BUB-1 and AIR-2 co-localise in the midbivalent ring domain during metaphase and their co-localisation decreases as anaphase progresses. Ultimately, BUB-1 is gone from the spindle and the bulk of AIR-2 is present in the central-spindle. **B.** BUB-1 and AIR-2 localisation was followed during meiosis I in oocytes expressing BUB-1::mCherry and GFP::AIR-2. Note that after kinetochore disassembly. **C.** BUB-1 and SUMO localisation was followed during meiosis I in oocytes expressing BUB-1::mCherry and GFP::SUMO. Scale bars, 2 µm.

In agreement with our previous data, BUB-1 and SUMO colocalise in the ring domain but no SUMO is detected in kinetochores during metaphase I (Figure 2C and Supplementary Movie 2). As anaphase progresses and kinetochores disassemble, BUB-1 kinetochore signal disappears, and BUB-1 concentrates in the stretched ring domains, as shown in fixed samples. At this stage, BUB-1 and SUMO display identical localisation patterns (Figure 2C and Supplementary Movie 2). During late anaphase, BUB-1 and SUMO not only display identical localisation but also behaviour: both protein become diffuse as they also decrease in intensity, until both proteins cease to be detected within the spindle (Figure 2C and Supplementary Movie 2). Therefore, SUMO and BUB-1 localise within the segregating chromosomes with their levels peaking at early anaphase and then both proteins leave the spindle during late anaphase.

### BUB-1 is a SUMO substrate and its localisation is regulated by SUMO

Given the striking BUB-1 and SUMO colocalisation observed during anaphase, we wondered whether BUB-1 could be conjugated by SUMO. We performed in vitro SUMO conjugation assays and determined that BUB-1 could be modified by SUMO and this modification increased with increasing amounts of the E2 conjugating enzyme UBC-9 (Figure 3A). Since high UBC-9 concentrations can lead to modification of otherwise unmodified lysines, we used limiting UBC-9 concentrations and increasing amounts of the meiotic SUMO E3 ligase GEI-17. BUB-1 SUMO modification is increased by GEI-17 in a dose-dependent manner (Figure 3B). We have shown before that depletion of SUMO or GEI-17 leads to the loss of BUB-1 from the midbivalent but not from kinetochores during metaphase of meiosis I (Pelisch et al., 2017). Since these previous results were obtained from fixed samples and were restricted to metaphase, we followed BUB-1 localisation in live oocytes expressing expressing endogenous GFP-tagged BUB-1. In agreement with our previous results with fixed samples (Pelisch et al., 2017), depletion of SUMO leads to a selective disappearance of BUB-1 from the midbivalent (Figure 3C, D). As anaphase progresses and kinetochores disassemble, this effect becomes more evident. Under normal conditions BUB-1 localises only in the stretched ring domains and depletion of SUMO leads to the complete absence of BUB-1 from the spindle (Figure 3C and Supplementary Movie 3). Similar results were obtained after depletion of the SUMO E3 ligase GEI-17 (Figure 3C, D). Quantification of BUB-1 localisation in the region between segregating chromosomes as anaphase progresses is depicted in Figure 3D. In line with these results, SUMO depletion completely abolishes MDF-1^*M ad*1^ localisation during anaphase I (Supplementary Figure 1). BUB-1 depletion is known to produce chromosome segregation defects, namely lagging chromosomes (Dumont et al., 2010). In agreement with this, we observed lagging chromosomes in more than 80% of BUB-1-depleted oocytes (Figure 3E). Although less penetrant, depletion of SUMO leads to a similar phenotype with more than 20% of oocytes exhibiting lagging chromosomes (Figure 3E). These results indicate that BUB-1 localisation is under strict control of a SUMO-dependent pathway and suggest that either SUMO partially regulates midbivalent BUB-1 function or that the midbivalent BUB-1 population is not the sole responsible of BUB-1 function during meiosis, with kinetochore BUB-1 being important as well.

**Fig. 3.**
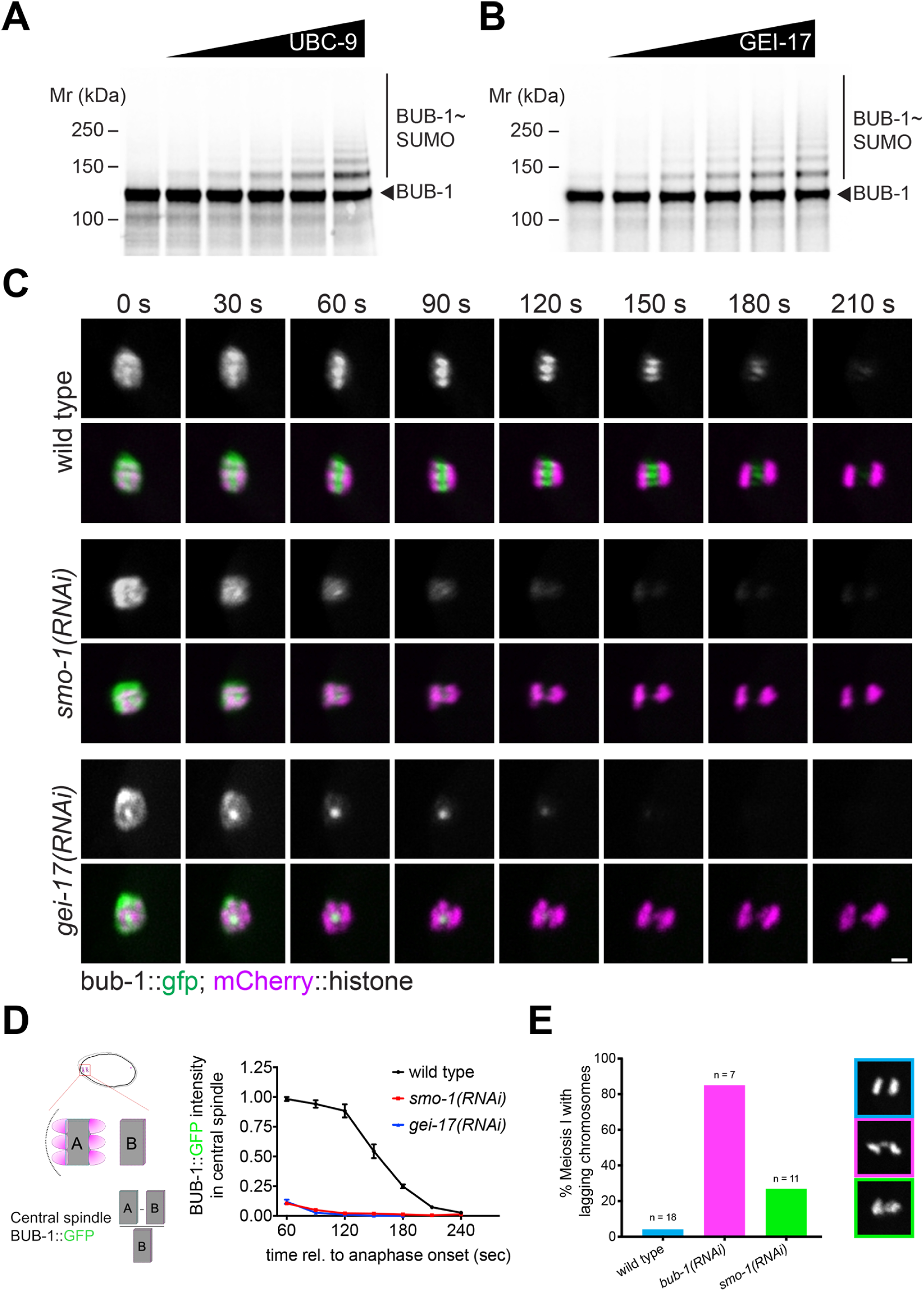
BUB-1 is a SUMO substrate and its localisation is regulated by SUMO. **A.** BUB-1 was incubated with increasing amount of UBC-9 and SUMO modification was analysed by SDS-PAGE. **B.** BUB-1 was incubated with a limiting amount of UBC-9 and increasing concentrations of the SUMO E3 ligase GEI-17 and the resulting reactions were analysed by SDS-PAGE to assess BUB-1 sumoylation. **C.** SUMO or GEI-17 were depleted by RNAi and BUB-1 localisation of was followed in a strain expressing BUB-1::GFP from the endogenous locus. Scale bar, 2 µm. **D.** Quantitation of the central-spindle BUB-1::GFP signal during anaphase. **E.** The graph shows the percentage of oocytes with lagging chromosomes during anaphase I after depletion of BUB-1 or SUMO.

### ULP-1 is an active SUMO protease in vivo and in vitro

Prompted by the sharp decrease in SUMO intensity during later anaphase, we thought to identify the SUMO protease(s) involved. In this line, a recent report has highlighted that ULP-1 plays a role during meiosis (Davis-Roca et al., 2018). To analyse SUMO protease activity in vivo, we used embryo extracts from wild type or ULP-1-depleted worms. We used embryos expressing GFP-tagged endogenous GEI-17, since ubiquitin and SUMO E3 ligases are known to undergo self-modification. GFP::GEI-17 was immunoprecipitated from extracts using an anti-GFP nanobody and autosumoylation was readily detected (Figure 4A). The identity of the slower migrating GFP::GEI-17 species was confirmed to contain SUMO using a SUMO specific antibody (Figure 4A, right blot). Upon depletion of ULP-1 by RNAi, we detected a large shift towards higher molecular weight forms of SUMO-modified GFP::GEI-17, leading to a complete disappearance of unmodified GFP::GEI-17 (Figure 4A). Therefore, ULP-1 has SUMO deconjugating activity in vivo. We then generated recombinant GEI-17 modified with fluorescently-labelled SUMO and incubated it with increasing amounts of full-length-ULP-1, generated by in vitro translation. In line with the in vivo results from Figure 4A, ULP-1 lead to a dose-dependent reduction in the amount of chains with a concomitant increase in the free SUMO (Figure 4B). While SUMO proteases can exhibit isopeptidase and peptidase activity, ULP-1 putative SUMO processing (peptidase) activity has not been tested to date. This is very important because results obtained after depletion of SUMO proteases that can perform both functions as the mammalian SENP1, could be more complicated to interpret. We then performed a SUMO processing assay using recombinant, unprocessed *C. elegans*. Since *C. elegans* SUMO has only one aminoacid after the SUMO C-terminal GlyGly motif, we added an HA tag to the C-terminus to allow for a better separation of the processed and unprocessed forms of SUMO after SDS-PAGE. Figure 4C shows that ULP-1 can process immature SUMO in a dose-dependent manner. Therefore ULP-1 can deconjugate SUMO from substrates and also process SUMO. Therefore, caution should be taken in depletion experiments because long depletions could in fact lead to depletion of the free, processed SUMO pool.

**Fig. 4.**
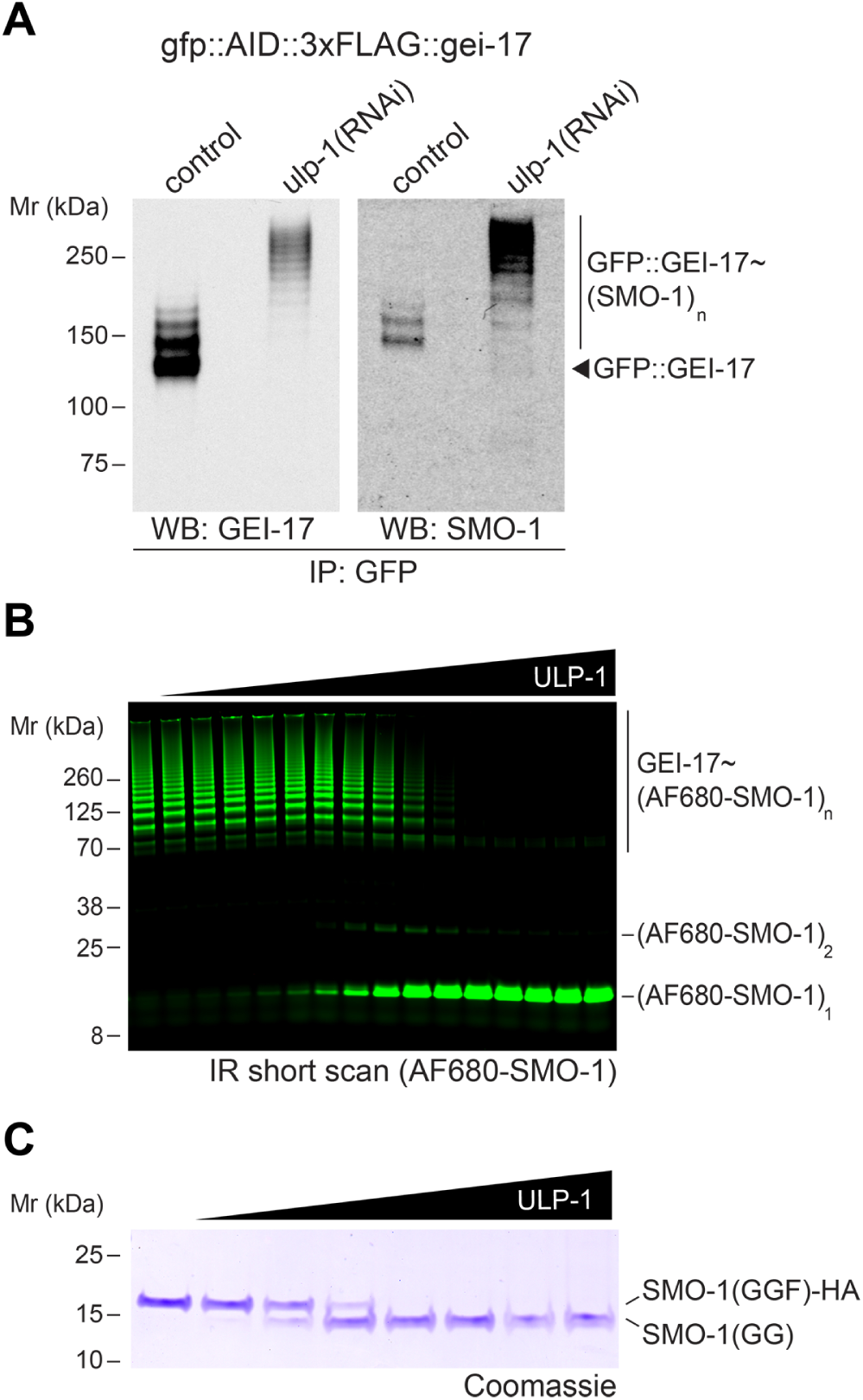
ULP-1 is an active SUMO protease in vivo and in vitro. **A.** Embryo extracts from wild type or *ulp-1(RNAi)*-fed worms were immunoprecipitated using an anti-GFP nanobody. The immunoprecipitate was analysed by western blot using GEI-17 and SUMO antibodies. **B.** Recombinant, Alexa Fluor 680-SUMO-modified GEI-17 was incubated with increasing amounts of ULP-1. The reactions were run on SDS-PAGE and scanned in a laser scanner. The identity of the different species on the gel are indicated on the right. **C.** Recombinant full length SMO-1-HA was incubated with increasing amounts of ULP-1 and the resulting reaction was run on SDS-PAGE and analysed by coomassie staining to resolve processed [‘SMO-1(GG)’] and unprocessed SUMO [‘SMO-1(GGF)-HA’].

### ULP-1 depletion leads to higher BUB-1 levels in the central spindle

Prompted by the finding of ULP-1 activity in vivo and in vitro, we went back to an in vitro setting and showed that ULP-1 can deconjugate SUMO from BUB-1, leading to completely unmodified BUB-1 (Figure 5A). We then used GFP-tagged endogenous ULP-1 to asses its localisation and no specific localisation was observed at any stage during meiosis I (Figure 5B and Supplementary Movie 4), while GFP::ULP-1 was readily detected throughout the mitotic spindle and nuclear envelope (Supplementary Figure 2 and Supplementary Movie 5). This result does not necessarily rule out a role for ULP-1 during meiosis because SUMO proteases are extremely active proteins and high concentrations would not be required for its activity in vivo. We then wondered whether ULP-1 affects BUB-1 localisation in vivo. In the absence of ULP-1, endogenous (GFP-tagged) BUB-1 displayed higher intensity throughout metaphase and anaphase (Figure 5C). Using fixed samples (and untagged BUB-1), we could also determine that it remains associated with the central spindle during later anaphase (Figure 5D). Furthermore, upon ULP-1 depletion, BUB-1 accumulates in foci and rod-like structures that co-localise with SUMO (Figure 5E). Therefore, ULP-1 is involved in regulating BUB-1 localisation during meiosis and its depletion has the opposite effect of depleting SUMO. The results also suggest that SUMO-modified BUB-1 is retained within the central spindle, by a yet to be characterised mechanism. We then analysed a putative role for ULP-1 during chromosome segregation. However, depletion of ULP-1 does not have a discernible effect on meiotic chromosome segregation (Figure 5F). It should be noted that a recent report found ULP-1 in the midbivalent and also attributed a more important role for ULP-1 during meiosis I (Davis-Roca et al., 2018). These results show that ULP-1 is an essential protein but the lethality is unlikely to arise due to meiotic defects. While we found no clear meiotic chromosome segregation defect, ULP-1 may still crucially regulate protein dynamics.

**Fig. 5.**
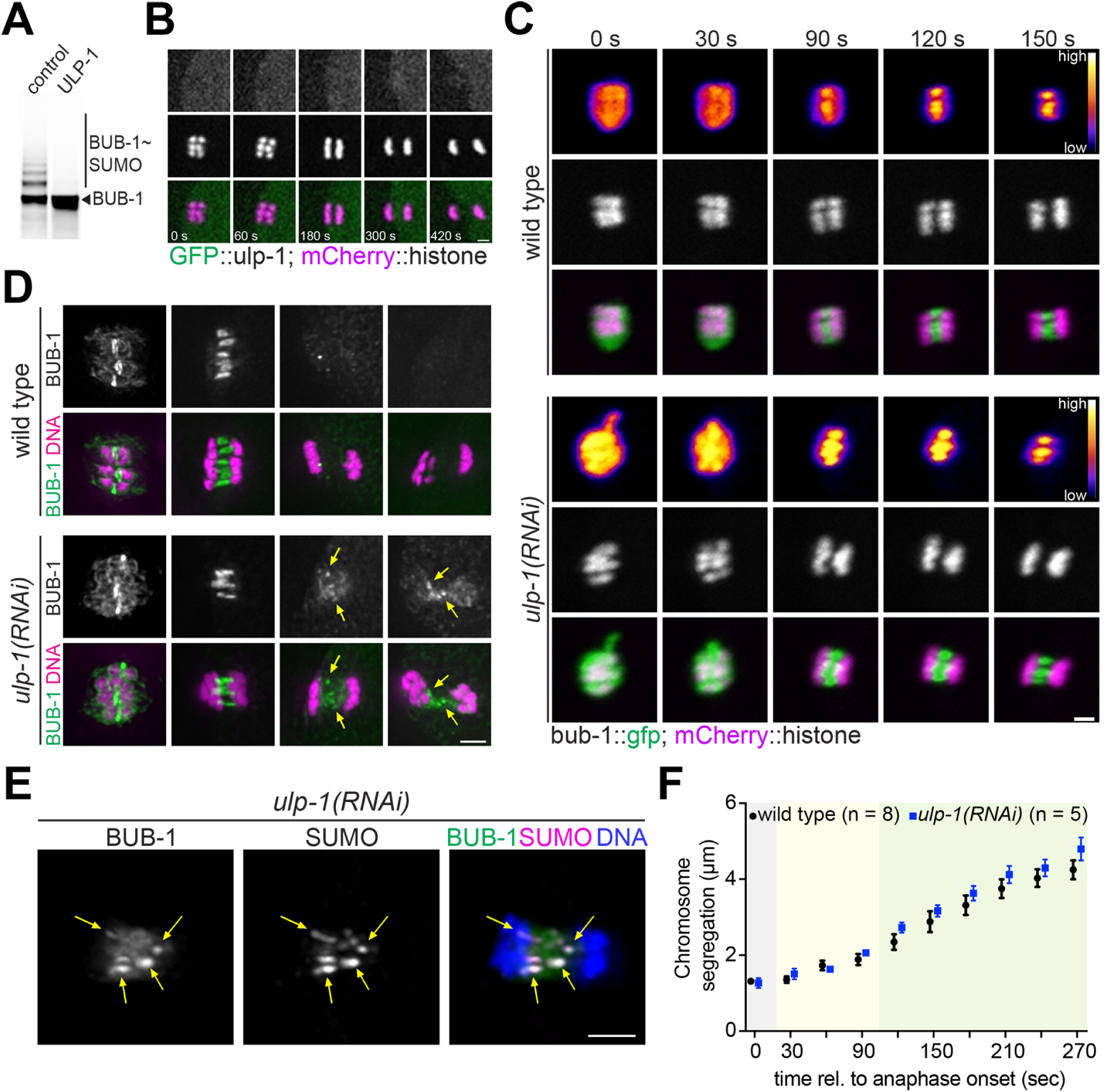
ULP-1 depletion and BUB-1 localisation. **A.** ULP-1 deconjugates SUMO from BUB-1. Full-length, in vitro-translated ULP-1 (or lysate control) were incubated with SUMO-modified BUB-1. **B.** Endogenous, GFP-tagged ULP-1 localisation was analysed during meiosis I in dissected oocytes. Scale bar, 2 µm. **C.** GFP-BUB-1 localisation I was followed during meiosis I in control (’wild type’) or ULP-1-depleted oocytes. Scale bar, 2 µm. The fire LUT scale runs from 3085 to 65000. **D.** BUB-1 localisation was assessed in fixed samples after ULP-1 depletion [*‘ulp-1(RNAi)*’] using a BUB-1-specific antibody. Scale bar, 2 µm. The yellow arrows indicate the foci where BUB-1 accumulates after ULP-1 depletion. **E.** BUB-1 and SUMO co-localisation during late anaphase after ULP-1 depletion. The yellow arrows indicate the aberrant accumulation of BUB-1 and its co-localisation with SUMO. Scale bar, 2 µm. **F.** Chromosome segregation was analysed in wild type and ULP-1-depleted oocytes.

### Central spindle CLS-2 localisation is regulated by SUMO

Another key protein during meiotic chromosome segregation is the CLASP orthologue CLS-2 (Dumont et al., 2010), whose presence in the central spindle is required for homologues to segregate during anaphase I (Dumont et al., 2010; Laband et al., 2017). We did not detect CLS-2 in the midbivalent ring domain during metaphase I or II (Supplementary Figure 3A). This difference is likely due to the appearance of kinetochore proteins on an end-on view of the spindle (Supplementary Figure 3A). In agreement with previous evidence (Laband et al., 2017), CLS-2 was detected in kinetochores and throughout the spindle during metaphase (Figure 6A and Supplementary Figure 3). During anaphase, CLS-2 was detected within the central-spindle (Figure 6A and Supplementary Figure 3B). Indeed, during anaphase, CLS-2 is detected more concentrated in areas close to the spindle-facing side of chromosomes (Supplementary Figure 3B, yellow arrows), which resemble the spots found for AIR-2 (Figure 2A, yellow arrows). When we depleted SUMO, we consistently observed a premature CLS-2 localisation within the midbivalent/central-spindle (Figure 6A and Supplementary Movie 6). SUMO-mediated regulation of CLS-2 seemed to be restricted to the early anaphase central spindle, since neither its kinetochore localisation or its late anaphase central-spindle localisation were affected (Figure 6A and Supplementary Movie 6). Depletion of the SUMO E3 ligase GEI-17 mirrored these results, reinforcing the notion that active sumoylation is taking place (Figure 6B). So far, our results are consistent with SUMO regulating early anaphase events, with an impact on BUB-1 and CLS-2 localisation.

**Fig. 6.**
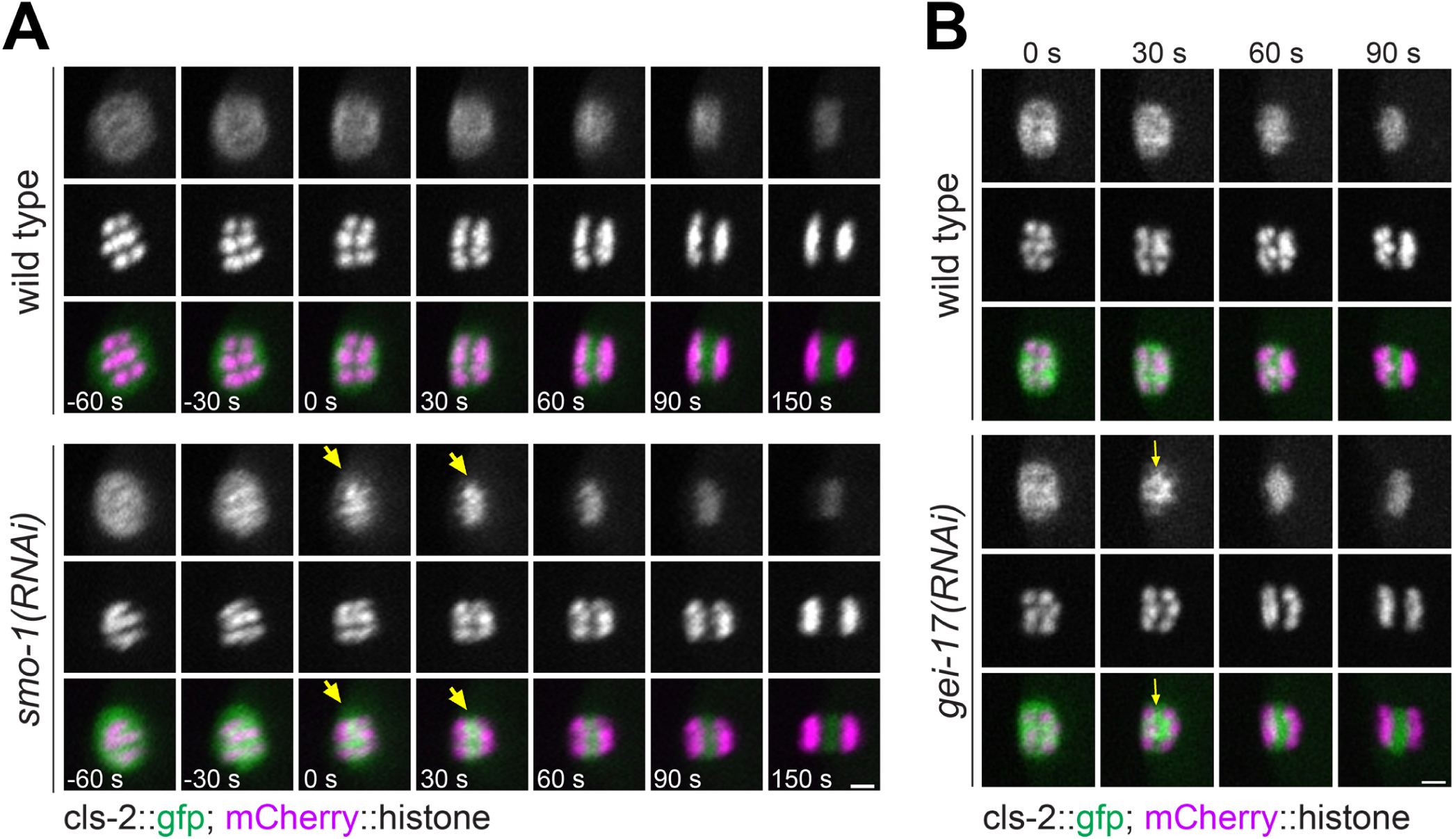
CLS-2 localisation is regulated by SUMO during early anaphase. **A.** CLS-2::GFP and mCherry::histone were followed during anaphase I in wild type and SUMO-depleted oocytes. The yellow arrows in the *smo-1(RNAi)* treated oocytes indicate the premature midbivalent/central-spindle CLS-2 localisation. **B.** Same as in (A), but depleting the SUMO E3 ligase GEI-17 instead of SUMO. Scale bars, 2 µm.

### Acute depletion of BUB-1 and CLS-2 during oocyte meiosis

The role of BUB-1 and CLS-2, as well as other proteins, has been mainly addressed using protein depletion by means of RNAi, either on its own or in depletion/rescue experiments (Davis-Roca et al., 2018; Dumont et al., 2010; Laband et al., 2017; Muscat et al., 2015; Wignall and Villeneuve, 2009). These experiments use RNAi mostly for between 12 and 48 hours, but in some cases RNAi incubation have reached up to 72 hours. This raises the concern that any identified phenotype during chromosome segregation after RNAi treatment could be, at least partially, due to defects in earlier meiotic events. We therefore used the auxin-induced degradation system to achieve acute protein depletion (Zhang et al., 2015) and conclusively rule out any potential early meiotic roles for BUB-1, and CLS-2. We generated strains carrying a fluorescent tag together with an auxin-inducible degron (AID) at their endogenous loci using CRISPR/Cas9 (Zhang et al., 2015). These strains also express fluorescently labelled histone as well as unlabelled TIR1 expressed only in germline and early embryos. Acute depletion of BUB-1 leads to defects in chromosome congression/alignment and segregation (Figure 7A and Supplementary Movie 7). as with RNAi-mediated BUB-1 depletion, chromosome segregation still took place, leading to the formation of the first polar body. We noted that the biggest effect of BUB-1 depletion occurred during Metaphase and early anaphase (Figure 7A and Supplementary Movie 7). Acute depletion of CLS-2 completely prevented chromosome segregation, and no polar body formation was observed under this conditions (Figure 7B and Supplementary Movie 8). Chromosome congression appeared normal in CLS-2-depleted oocytes (Supplementary Movie 9). We noticed that in some oocytes, chromosomes did start to segregate, although with severe defects, with meiosis I failing to extrude a polar body (Figure 7B and Supplementary Movie 9). Altogether, these results show that BUB-1 and CLS-2 play specific, non-overlapping roles during meiotic chromosome segregation.

**Fig. 7.**
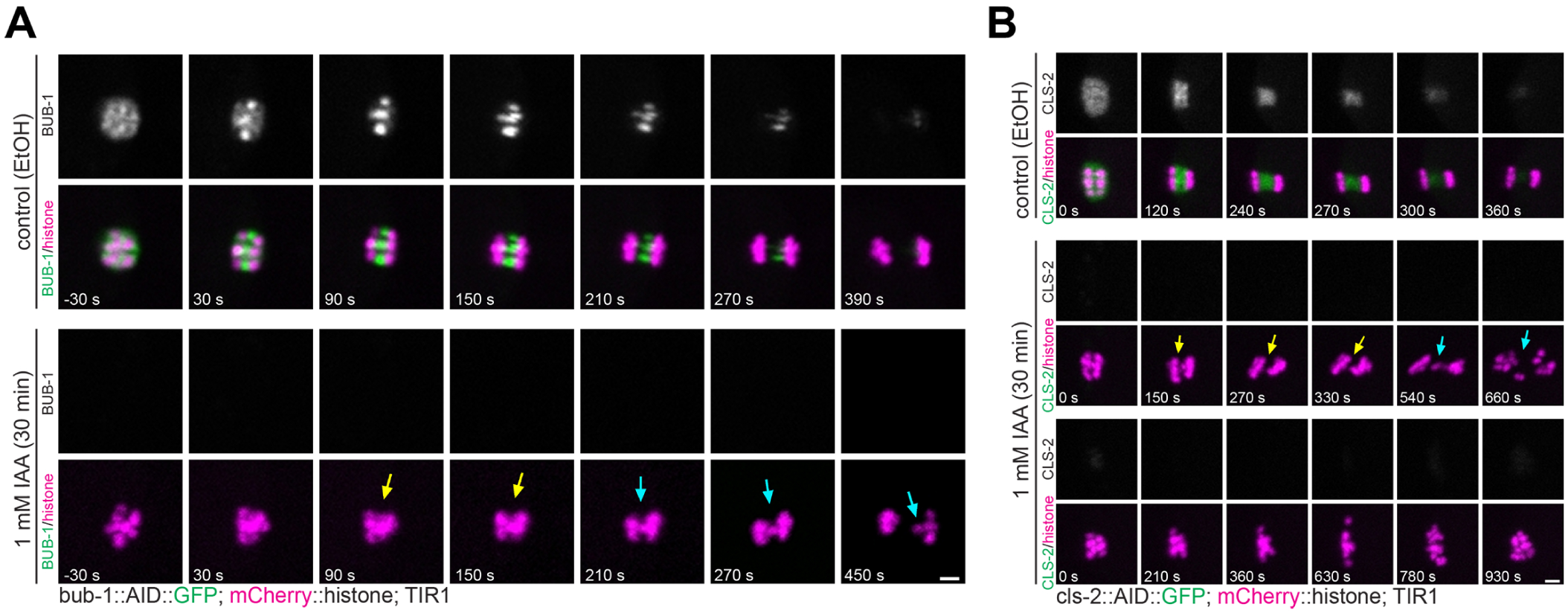
Acute depletion of BUB-1 and CLS-2. **A.** Endogenous BUB-1 was tagged with an auxin-inducible degron (AID) and GFP. Worms expressing untagged TIR1 were either treated with vehicle (ethanol) or auxin (‘IAA’), dissected, and oocytes were imaged. The yellow arrows highlight the early anaphase chromosome segregation defect observed after BUB-1 depletion, while the cyan arrows mark the lagging chromosomes during midand late anaphase. Scale bar, 2 µm. **B.** Endogenous CLS-2 was tagged with an auxin-inducible degron (AID) and GFP and its localisation and effects of its depletion were analysed as in (A). We noted that in some cases, chromosomes achieve some degree of separation after acute CLS-2 depletion (yellow arrows), however in all cases, segregation failed and no polar bodies were extruded. Scale bar, 2 µm.

**Fig. 8.**
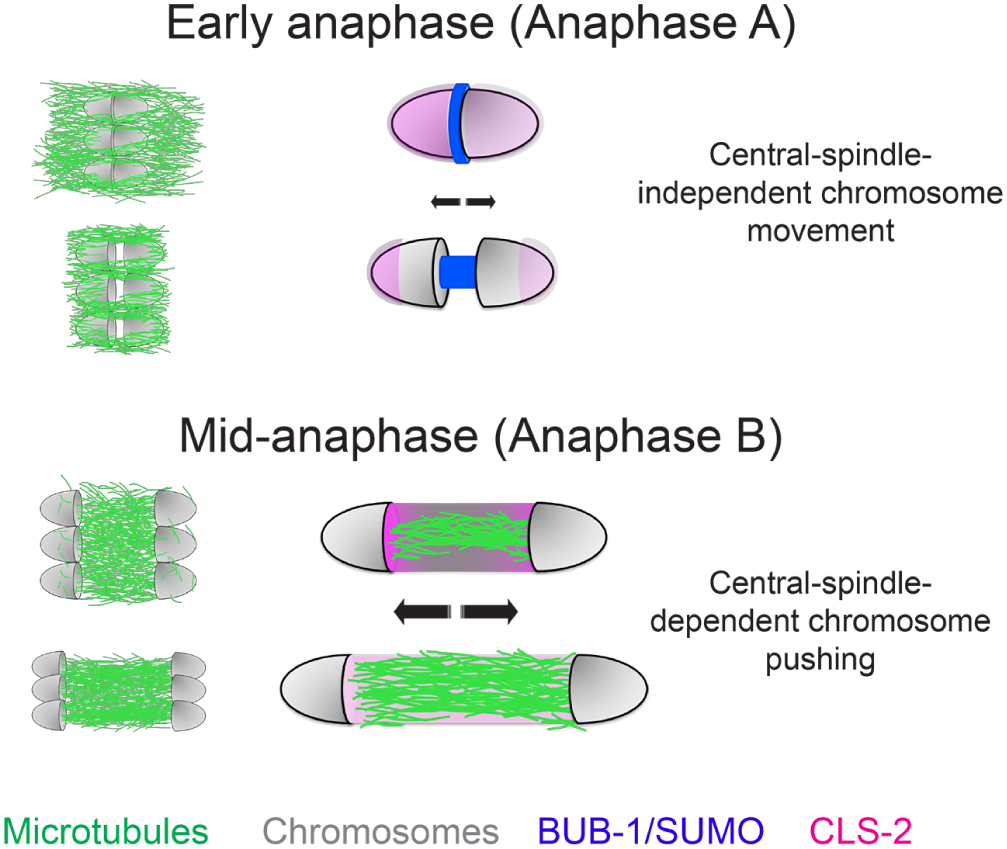
Two-step chromosome segregation model and the role of SUMO. During early anaphase, chromosomes begin to separate without microtubules being present between them. This area is filled with BUB-1 and SUMO, among other proteins, suggesting that these proteins could play a role during this early segregation step. As anaphase progresses, microtubules populate the region between segregating chromosomes leading to the CLS-2-dependent stage.

## Discussion

Here, we show that SUMO modification regulates central spindle protein localisation. We found clear defects in the localisation of the spindle checkpoint components BUB-1 and MDF-1, and the CLASP orthologue CLS-2. Midbivalent ring domain BUB-1 is subject to control by the SUMO E3 ligase GEI-17 and the SUMO protease ULP-1. In contrast, kinetochore BUB-1 is unaffected by the SUMO-mediated control. In addition, we have shown that BUB-1 is a SUMO substrate and its modification is determined by GEI-17-mediated conjugation and ULP-1-mediated deconjugation. Altogether, we propose sumoylation as an emerging PTM required for the tight spatial and temporal regulation of proteins involved in oocyte chromosome segregation.

### SUMO-dependent regulation of BUB-1 and CLS-2 localisation

It has become increasingly clear that the midbivalent ring domain does not behave as a ‘static’ entity. Its composition changes dramatically during metaphase-anaphase I and each protein displays a characteristic and dynamic localisation pattern (Wignall and Villeneuve, 2009; Dumont et al., 2010; Collette et al., 2011; Connolly et al., 2015; Han et al., 2015.; Muscat et al., 2015; Gigant et al., 2017). While SUMO depletion on its own does not drastically affect chromosomes segregation, it regulates the dynamic localisation of midbivalent/central-spindle proteins. During metaphase I, BUB-1 localisation in the midbivalent is strictly dependent on SUMO conjugation (Pelisch et al., 2017). During anaphase, kinetochores disassemble and BUB-1 is concentrated in rod-like structures in the central-spindle, and this localisation is also entirely dependent on SUMO conjugation.

While CLS-2 plays a key role in chromosome segregation, regulators of its activity and or/localisation have not been characterised. While kinetochore localisation of CLS-2 depends entirely on BUB-1 (Dumont et al., 2010; Laband et al., 2017), central spindle localised CLS-2 is detected after BUB-1 depletion (Laband et al., 2017). Our results show that timely CLS-2 localisation in the midbivalent/central spindle depends on SUMO: after SUMO depletion, CLS-2 appears to leave kinetochores and concentrate between the homologous chromosomes prematurely (Figure 3C). This raises the intriguing possibility that different BUB-1 populations could regulate CLS-2 in different ways. In this scenario, kinetochore-localised BUB-1 would positively regulate CLS2 localisation while midbivalent BUB-1 would inhibit CLS-2 localisation. Still in the speculative arena, SUMO could be a switch for this dual behaviour displayed by BUB-1. SUMO-modified and/or SUMO-bound BUB-1 could lead to a disruption in its interaction with CLS-2, which likely occurs via the CENP-F orthologues HCP-1 and HCP-2 (Dumont et al., 2010). Other factors could certainly be involved in this regulation and further experiments are required to test this model.

### CLS-2^*CLASP*^, and BUB-1^*Bub*1^ roles in meiotic chromosome segregation are not due to early meiotic events

The architecture of the *C. elegans* germline has been one of the key advantages of this model system allowing for very efficient mRNA depletion via RNAi. However, when focusing on chromosome segregation, this could lead to erroneous interpretations, in particular for experiments utilising long RNAi incubations. In those cases, depletion can have an impact on early meiotic events such that chromosome segregation is affected partly or even solely by this alteration of these events. Therefore, acute inactivation or depletion at the protein level is required to analyse chromosome segregation independently of previous meiotic events. One possibility is the use of fast acting temperature-sensitive alleles (Severson et al., 2000; Davies et al., 2014). We used tissuespecific, auxin-induced degradation in worms, as introduced by the Dernburg lab (Zhang et al., 2015). We could determine that protein depletion was achieved in most cases in under one hour. Acute depletion of the CPC component AIR-2 (data not shown) or the CLASP orthologue CLS-2 (Figure 7B) lead to a complete failure in chromosome segregation. We did notice two different scenarios: i) chromosomes completely failed to separate from anaphase onset and ii) chromosomes were able to initiate separation, but later failed to achieve proper segregation and polar body extrusion (Figure 7B). While future experiments will determine the cause of these two different phenotypes, we speculate that CLS-2 might not be essential for the very first steps in chromosome separation during anaphase. This notion would go in line with the fact that MTs populate the area between segregating chromosomes later during anaphase (Redemann et al., 2018), and it is then when a more relevant role for CLS-2 would take place. In the case of BUB-1, acute depletion lead to a similar phenotype to that of RNAi-mediated depletion (Dumont et al., 2010): defective segregation indicated by the presence of lagging chromosomes. As opposed to AIR-2 and CLS-2 depletions, chromosomes did segregate and polar bodies were formed, suggesting that AIR-2 is more likely than BUB-1 to be a key CLS-2 regulator during anaphase. This is also supported by the presence of both AIR-2 and CLS-2 in foci next to chromosomes during early anaphase (Figure 2A and Supplementary Figure 3B). In sum, AIR-2 and CLS-2, and BUB1 could play different and partially overlapping roles during meiotic chromosome segregation and these roles are not due to early meiotic defects. We envision that the AID system will provide more accurate interpretations in the future, also in the study of the first mitotic divisions, to avoid the impact of known or unknown meiotic defects.

### How is initial chromosome separation achieved during anaphase

Several lines of evidence point towards a two-step chromosome segregation mechanism operating during anaphase I in *C. elegans* oocytes: i) Two phases of chromosome segregation characterised by different segregation speeds were reported (McNally et al., 2016); ii) central-spindle CLS-2 localisation is at least partially BUB1-independent (Laband et al., 2017); iii) central-spindle ablation during mid-anaphase stops chromosomes segregation (Laband et al., 2017; Yu et al., 2019); and iv) the area between segregating homologues is microtubule-free during early anaphase (Redemann et al., 2018), suggesting that the initial steps of segregation could be at least partially CLS2-independent. Our results suggest that this early anaphase stage is subject to regulation by SUMO, and it will therefore be important to address which protein(s) are downstream of BUB-1 and SUMO. Some interesting targets of the initial chromosome movement are motor proteins. In particular, we have observed that the CENP-F orthologue HCP-1 has an interesting localisation pattern: while its localisation during metaphase mirrors that of CLS-2 and it is also under the control of BUB-1, HCP-1 populates the midbivalent or central-spindle region earlier than CLS-2, and also concentrates in spots close to DNA, where the CPC and CLS-2 are (data not shown). Therefore, HCP-1 (and its paralogue HCP-2) could be regulating events during early anaphase, in addition to recruiting CLS-2 during mid-anaphase. While more experiments are needed to test this hypothesis, an intricate interaction between BUB-1, HCP-1/2, and CLS-2 has recently been involved in the regulating kinetochore-MT attachments during mitosis (Edwards et al., 2018).

### The SUMO protease ULP-1

A recent paper has found that ULP-1 depletion does have a more dramatic impact on chromosome segregation (Davis-Roca et al., 2018). A possible explanation for this discrepancy is the long RNAi incubations used in the mentioned study. We did find that long incubation with *ulp-1(RNAi)* has a more dramatic effect, mimicking that of SUMO depletion. Indeed, since we report that ULP-1 is a SUMO processing enzyme, long depletions are likely to lead to the absence of mature SUMO available for conjugation. In spite of this discrepancy, desumoylation does have a role at least in regulating protein dynamics within segregating chromosomes and more experiments are needed to shed light into the underlying mechanisms.

### Concluding remarks

Overall, we have shown that sumoylation regulates the dynamics of central-spindle proteins during female meiosis, namely BUB-1 and CLS-2. Previous reports have highlighted the importance of the central-spindle for chromosome segregation in oocytes (Laband et al., 2017; Redemann et al., 2018; Yu et al., 2019). Remarkably, this central-spindle-based mechanism could be more widespread than anticipated, as it has been shown to exist also during mitosis in *C. elegans* and in human cells (Yu et al., 2019). Here we focused on the dynamic behaviour of key proteins and to what extent this is regulated by sumoylation. Our findings show that precise dynamic localisation of the kinase BUB-1 and the CLASP orthologue CLS-2 is dependent on sumoylation. It is important to note that this is likely not to be the only mechanism regulating these proteins’ localisation since depletion of either SUMO or the SUMO protease ULP-1 do not have a drastic effect during chromosome segregation. In spite of this, and given the increasing relevance of the central-spindle during early anaphase, understanding how protein function and localisation within the central-spindle are regulated will be key in obtaining the full picture of the different mechanisms driving chromosome segregation. In this context, PTMs such as sumoylation and phosphorylation are likely to play fundamental roles during chromosome segregation. Interestingly, we have shown that meiotic phosphorylation in the midbivalent is dependent on SUMO and these two PTMs acting together would contribute a great degree of versatility to the system (Pelisch et al., 2017). Future studies will provide insight into this and probably other PTM crosstalk taking place during cell division.

## Methods

### *C. elegans* strains

Strains used in this study were maintained at 20 degrees unless unless indicated otherwise. For a complete list of strains, please refer to Table 1. Requests for strains not deposited in the CGC should be done through FP lab’s website (https://pelischlab.co.uk/reagents/).

**Table 1.**
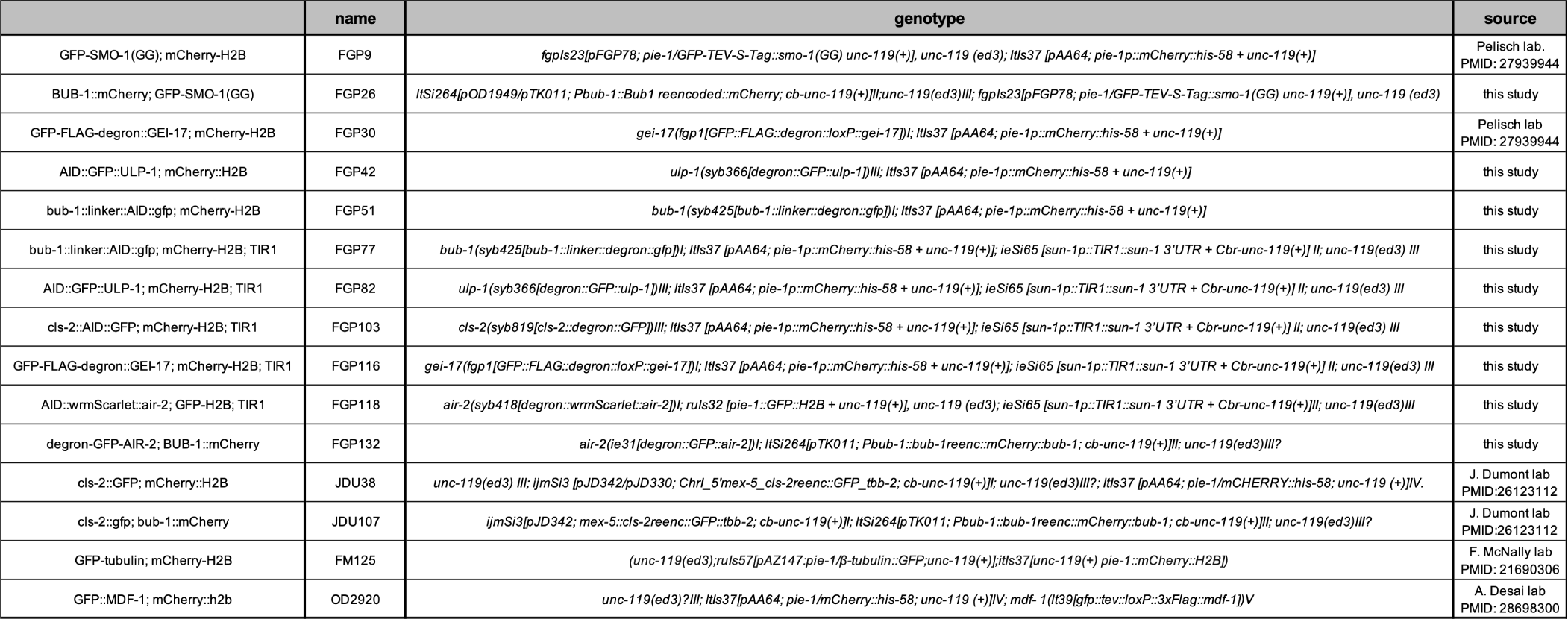
*C. elegans* strains used in this study

### Auxin-induced protein degradation

All the germline-expressing TIR1 strains were generated by the Dernburg lab (Zhang et al., 2015). For live imaging, we used the strain CA1353 as it contains an untagged version of TIR1. The degron sequence used in this study consisted of the 44-aa fragment of the Arabidopsis thaliana IAA17 protein (Morawska and Ulrich, 2013; Zhang et al., 2015). Auxin was used at 1 mM final concentration in standard NGM plates, unless otherwise noted. All plates for auxin treatment were prepared, allowed to dry for 2 days and a lawn of concentrated OP50 bacteria was seeded, as auxin inhibits bacterial growth. For auxin treatment, worms were placed on auxin-containing plates for the indicated times.

### Live imaging of oocytes

A detailed protocol for live imaging of *C. elegans* oocytes was used with minor modifications (Laband et al., 2018). Fertilized oocytes were dissected and mounted in 5 µl of L-15 blastomere culture medium (0.5 mg/mL Inulin; 25 mM HEPES, pH 7.5 in 60% Leibowitz L-15 medium and 20% heat-Inactivated FBS) on 24×40 mm coverslips. Once dissection was performed and early oocytes identified using a steremicroscope, a circle of vaseline was laid around the sample, and a custom-made 24X40 mm plastic holder (with a centered window) was placed on top. The sample was immediately transferred for imaging. Live imaging was done using a 60X/NA 1.4 oil objective on a spinningdisk confocal microscope (MAG Biosystems) mounted on a microscope (IX81; Olympus), a Cascade II camera (Photometrics), spinning-disk head (CSU-X1; Yokogawa Electric Corporation). Acquisition parameters were controlled by MetaMorph 7 software (Molecular Devices). For all live imaging experiments, maximal projections are presented. Figures from live imaging experiments were prepared using OMERO.figure.

### Immunofluorescence

Worms were placed on 4 µl of M9 worm buffer in a poly-D-lysine (Sigma, P1024)-coated slide and a 24 × 24-cm coverslip was gently laid on top. Once the worms extruded the embryos, slides were placed on a metal block on dry ice for >10 min. The coverslip was then flicked off with a scalpel blade, and the samples were fixed in methanol at 20°C for 30 min (except for GFP, where the methanol treatment lasted 5 min). Embryos were stained using standard procedures. Secondary antibodies were donkey anti–sheep, goat anti-mouse, or goat anti-rabbit conjugated to Alexa Fluor™ 488, Alexa Fluor™ 594, and Alexa Fluor™ 647 (1:1,000, Thermo Scientific). Donkey anti-mouse and donkey anti-rabbit secondary antibodies were obtained from Jackson ImmunoReserach. Embryos were mounted in ProLong Diamond antifade mountant (Thermo Scientific) with DAPI.

### GFP immunoprecipitation

For GFP immunoprecipitations, we followed a published protocol (Sonneville et al., 2017) with minor modifications. Approximately 1000 worms expressing GFP-tagged endogenous GEI-17 were grown for two generations at 20°C in large 15-cm NGM plates with concentrated HT115 bacteria. Worms were bleached and embryos were laid in new 15-cm NGM plates with concentrated HT115 bacteria. Once at the L4 stage, worms were washed and placed on 15-cm agarose plates containing concentrated *ulp-1(RNAi* or empty L4440 vector transformed bacteria. After 25 hs, worms were bleached and the embryos were resuspended in a lysis buffer containing 100 mM HEPES-KOH pH 7.9, 50 mM potassium acetate, 10 mM magnesium acetate, 2 mM EDTA, 1X Protease inhibitor ULTRA (Roche), 2X PhosSTOP (Roche), 1 mM DTT, and 10 mM iodoacetamide. The solution was added drop-wise to liquid nitrogen to generate beads that were later grinded using a SPEX SamplePrep 6780 Freezer/Mill. After thawing, we added one-quarter volume of buffer containing lysis buffer supplemented with 50% glycerol, 300 mM potassium acetate, 0.5% NP40, plus DTT and protease and phosphatase inhibitors as above. DNA was digested with 1,600U of Pierce Universal Nuclease for 30 min on ice. Extracts were centrifuged at 25,000 g for 30 min and then at 100,000 g for 1 h. The extract was then incubated for 60 min with 30 µl of a GFP nanobody covalently coupled to magnetic beads. The beads were washed ten times with 1 ml of wash buffer (100 mM HEPES-KOH pH 7.9, 300 mM potassium acetate, 10 mM magnesium acetate, 2 mM EDTA, 0.1% NP40, plus protease and phosphatase inhibitors). Bound proteins were eluted twice using 50 µl LDS sample buffer (Thermo) at 70 °C for 15 min and stored at −80°C.

### Antibody labelling

For all experiments involving fluorescence intensity measurements, antibodies were labelled with Alexa fluorophores. The APEX Alexa Fluor labelling kits (Thermo Scientific) were used and antibodies were labelled with Alexa Fluor™ 488, Alexa Fluor™ 594, and Alexa Fluor™ 647, following the manufacturer’s indications. Antibodies were buffer exchanged to PBS using Zeba™ Spin De-salting Columns (Thermo Scientific) and were stored in small aliquots at −20°C in PBS containing 10% glycerol. Labelled antibodies were used at 1-5 µg/ml for immunofluorescence.

### Protein production

Full-length BUB-1 and ULP-1 cDNAs were cloned into pF3 WG (BYDV) Flexi^®^ Vectors and expressed using the TnT® SP6 High-Yield Wheat Germ Protein Expression System (Promega). BUB-1 reactions included 35[S]-labelled methionine to allow for further detection, whereas ULP-1 reactions were left unlabelled. GEI-17, UBC-9, and all SUMO variants were expressed and purified as described previously (Pelisch et al., 2014; Pelisch et al., 2016; Pelisch et al., 2017). SUMO labelling was achieved using Alexa Fluor™ 680 C2 Maleimide (Thermo). Reactions were performed according to the manufacturer’s protocol. A cysteine residue at position 2 was created in *C. elegans* SUMO, leading to SMO-1(A2C). Untagged SMO-1(A2C) was purified using the same protocol used for wild type SMO-1. After labelling, we got rid of the free dye by gel filtration. The product was analysed by mass spectrometry and confirmed the absence of free dye. We checked that the mutant SUMO behave like the wild type in thioester formation as well multiple turnover conjugation reactions.

### GEI-17 autosumoylation

GEI-17 automodification was carried out in the following conditions: 50 mM Tris-HCl pH 7.5, 0.5 mM TCEP, 2 mM ATP, 5 mM *MgCl*_2_, 2 µM labelled SMO-1, 6 µM UBC-9, 6 ng/µl of human E1, and 0.5 µM GEI-17(133-509). SUMO-modified GEI-17 was further purified by size exclusion chromatography using a Superdex 200pg column. This step removed any free SUMO and SUMO-conjugated UBC-9 from the reaction.

### ULP-1 treatment of SUMO-modified GEI-17

SUMO deconjugation was performed by adding 1 µl of the ULP-1 expression reaction to 12.5 µl of SUMO-modified GEI-17 and incubating for 1 hour at 37°C.

### BUB-1 sumoylation and desumoylation

One µl of 35^*S*^ Methionine labelled BUB-1 was incubated with 60 ng of human SUMO E1, 500 ng UBC-9 (for E3-independent reactions) or 30 ng UBC-9 (for GEI-17-dependent reactions), 1 µg of SUMO per 10 µl. Reactions were performed in 50 mM Tris-HCl pH 7.5, 0.5 mM TCEP, 2 mM ATP, 5 mM *MgCl*2, 10 mM creatine phospohate, 3.5 U/ml creatine kinase, 0.6 U/ml inorganic pyrophosphatase, and 1X Protease inhibitor cocktail (cOmplete, Roche). reactions were incubated for 4 hs at 37°C. Samples were either analysed for SUMO conjugation or treated with ULP-1 before analysis. For ULP-1 treatment, 25 µl reactions were incubated with 1 µl of ULP-1 mix (or vector control) for 2 hs at 30*°*C.

### SUMO processing

SUMO processing was performed on an C-terminal HA-tagged version of full-length SMO-1 (Pelisch et al., 2014). three µg of SUMO were incubated with either 1 µl or 1 µl of serial 1/2 dilutions of the ULP-1 expression reaction in the presence of 50 mM Tris-HCl pH 7.5, 150 mM NaCl, and 0.5 mM TCEP. For all ULP-1 treatments, extracts with empty vector were used as a control. SUMO processing by ULP-1 was analysed by coomassie staining or by western blot using mouse anti-HA and sheep anti-SMO-1 antibodies. Blots and reaction containing Alexa Fluor™ 680 were analysed with an Amersham Typhoon 5 Biomolecular Imager.

## ACKNOWLEDGEMENTS

We would like to thank Abby Dernburg for sharing strains and advise on CRISPR and auxin-induced degradation protocols. We would also like to thank Arshad Desai for sharing strains and antibodies and Julien Dumont for sharing strains. We are grateful to Tomo Tanaka and Dhanya Cheerambathur for comments on the manuscript. We also thank Ricardo Henriques for making the overleaf template used for writing and formatting this manuscript publicly available. This work was supported by a Career Development Fellowship from the Medical Research Council (grant MR/R008574/1 to FP), an ISSF grant funded by the Wellcome Trust (105606/Z/14/Z to FP), and an Investigator Award from the Wellcome Trust (098391/Z/12/7 to R.T.H). Some nematode strains were provided by the CGC, which is funded by NIH Office of Research Infrastructure Programs (P40 OD010440). We thank the Dundee Imaging Facility, which is supported by the ‘Wellcome Trust Technology Platform’ award (097945/B/11/Z) and the ‘MRC Next Generation Optical Microscopy’ award (MR/K015869/1), and the Tissue Imaging Facility, funded by a Wellcome Trust award (101468/Z/13/Z).

