## Supplementary figures and images for "Sumoylation regulates central spindle protein dynamics during chromosome segregation in oocytes"

### Supplemental Figure 1

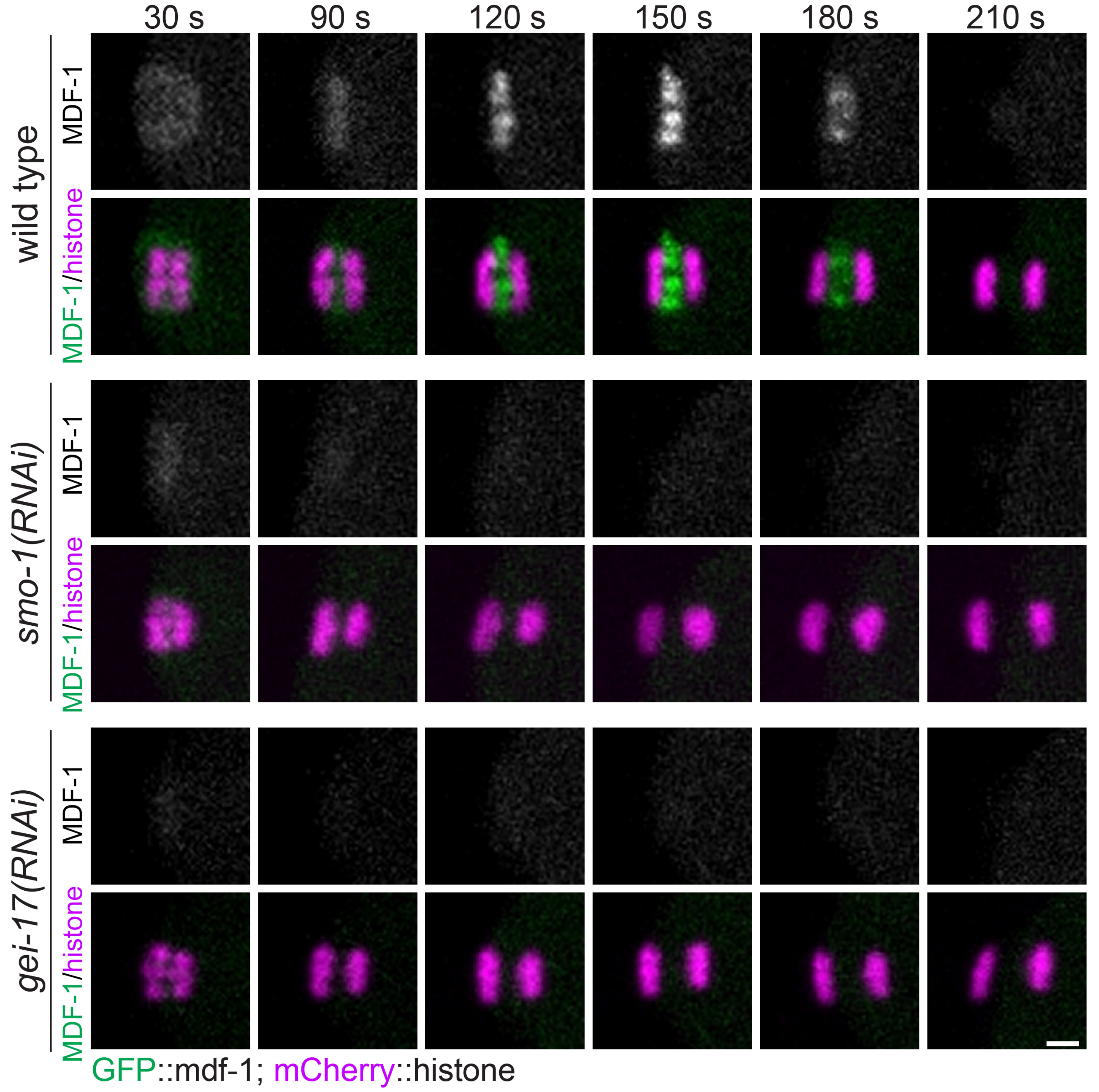

### Supplemental Figure 2

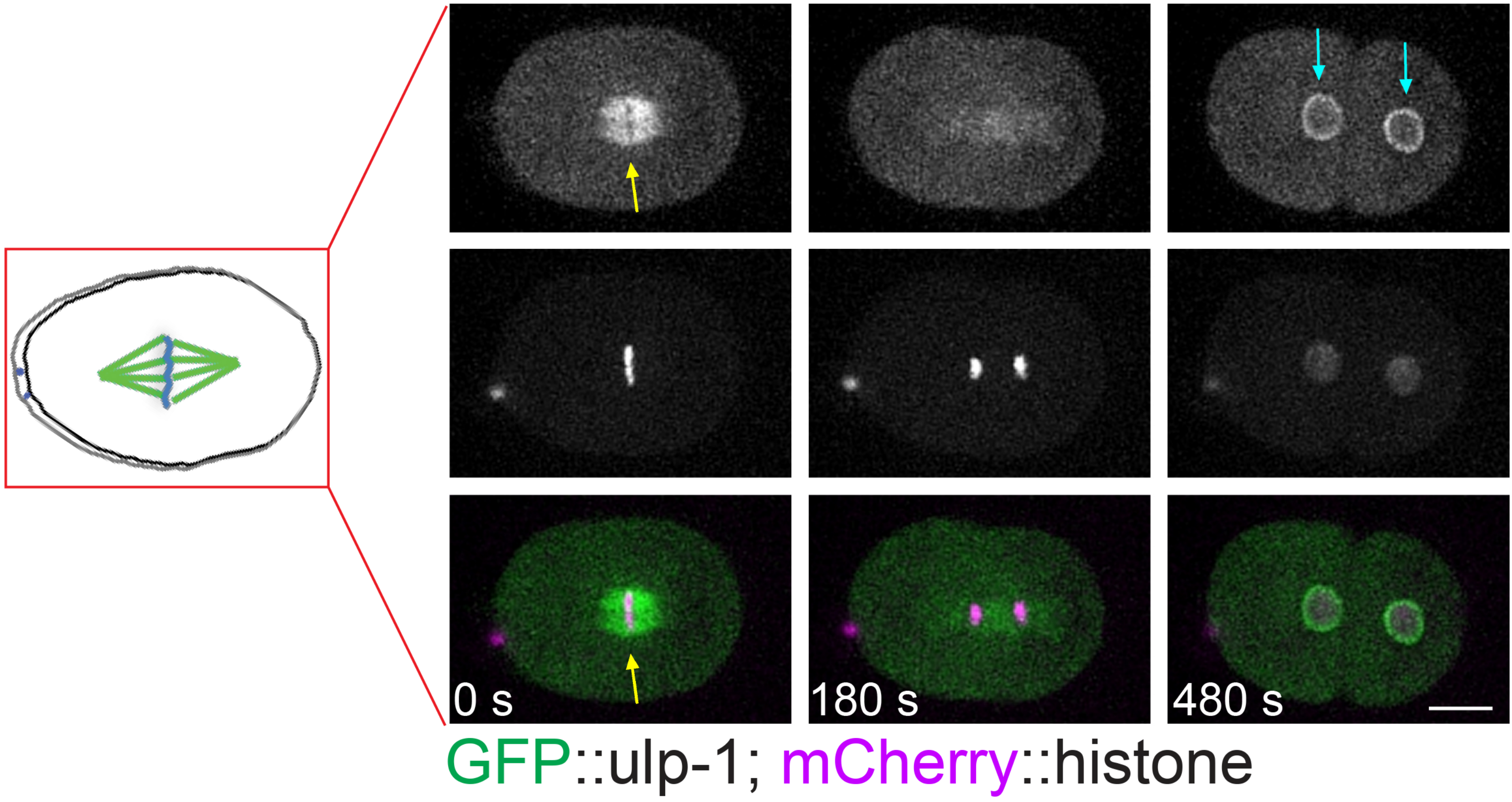

### Supplemental Figure 3

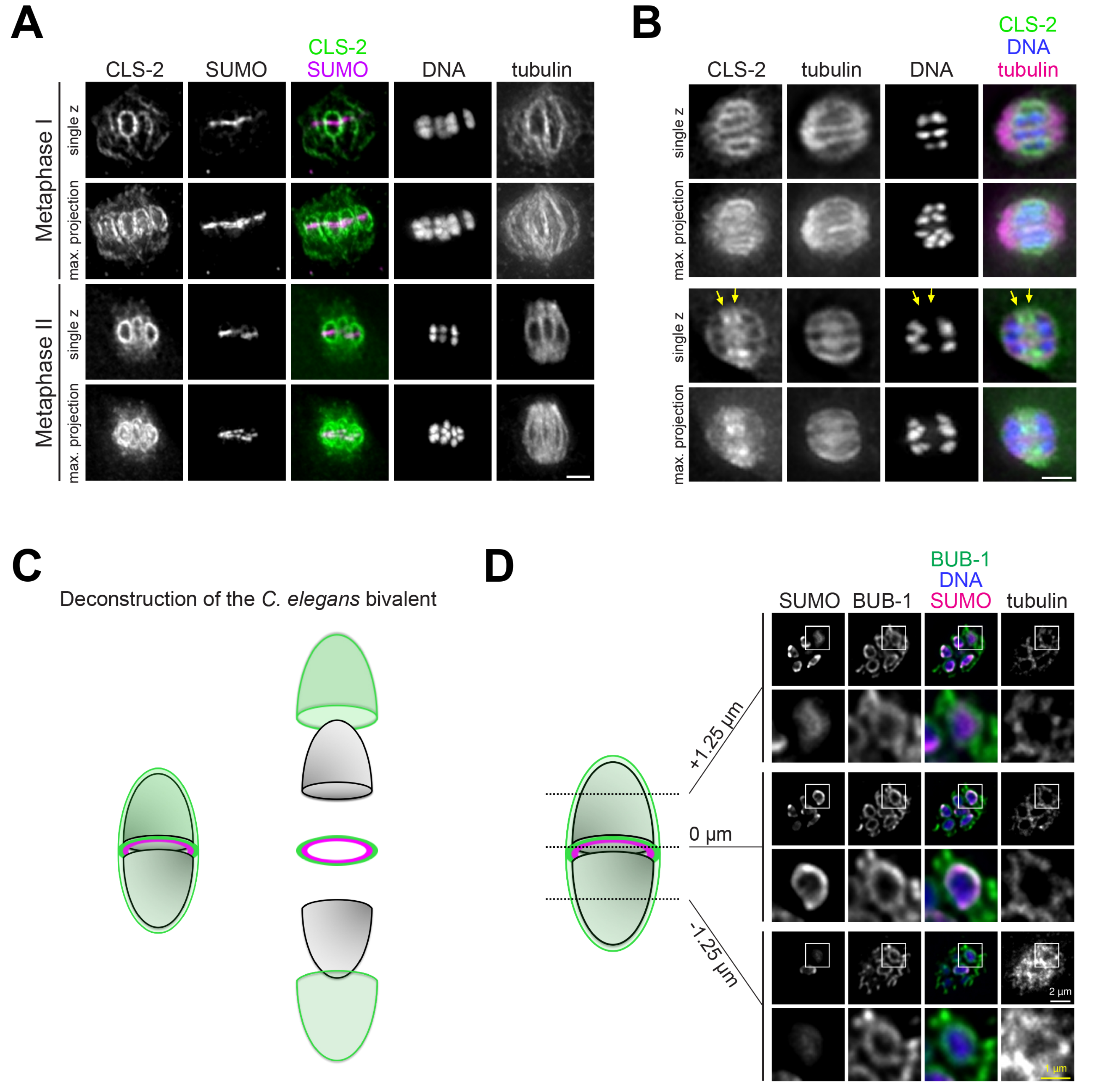
